# Null Subtraction Imaging for Functional Ultrasound Brain Activation Mapping

**DOI:** 10.64898/2026.04.14.718533

**Authors:** Gonzalo Garay, Juan Barolin, Victoria Sorriba, Juan Pablo Damián, Zhengchang Kou, Michael Oelze, Carlos Negreira, Alejandra Kun, Javier Brum

## Abstract

Null Subtraction Imaging (NSI) is a nonlinear beamforming approach that combines multiple receive apodizations and subtraction to improve spatial resolution in ultrasound imaging. In NSI, a DC offset parameter is introduced in the apodization design to control the sharpening of the effective beam pattern and, therefore, the degree of spatial-resolution enhancement. Here, we investigate the use of NSI in functional ultrasound (fUS) imaging of the mouse brain and compare its performance with conventional delay-and-sum (DAS) beamforming across a range of DC offset values. fUS acquisitions were performed in three anesthetized wild-type mice during periodic vibrissae stimulation. Activation maps were computed by correlating cerebral blood volume (CBV) signals with the stimulation pattern. Activation area, edge gradient, Dice similarity coefficient, and signal-to-noise ratio (SNR) were used to evaluate spatial localization, boundary sharpness, vascular alignment and signal stability, respectively. NSI yielded more spatially confined activation maps than DAS and produced sharper activation boundaries. However, for low DC offsets (*DC* < 0.5), the CBV signal exhibited increased fluctuations, which reduced temporal stability and limited the reliability of the functional maps. As the DC offset increased, temporal SNR improved, while the spatial-resolution gain progressively decreased. In our imaging configuration, intermediate DC values around *DC* ≈ 0.5 provided the most favorable compromise between improved spatial localization and sufficient temporal stability for reliable functional activation detection. These results demonstrate the feasibility of applying NSI to functional ultrasound imaging and provide a quantitative framework for selecting the DC parameter in fUS studies.

## 1. Introduction

In fUS, power Doppler images are typically acquired at temporal sampling rates between 1 and 10 Hz, enabling the assessment of brain function through CBV changes induced by neurovascular coupling. This provides an indirect measure of neuronal activity, both at rest and in response to external stimuli [1, 2, 3]. fUS has been successfully used to map the dynamics and connectivity of brain activity across different cognitive states [4] in rodents [5], non-human primates [6] and humans [7, 8, 9], establishing it as a versatile tool in modern neuroscience [10, 11, 12]. Despite these advantages, the spatial resolution of fUS remains limited by diffraction, which sets a resolution limit on the order of half the ultrasound wavelength. One possible way to improve spatial resolution is to increase the transmitted frequency, but this comes at the cost of reduced penetration depth due to higher attenuation.

As an alternative, several ultrasound methods have recently been developed to enhance spatial resolution without losing imaging depth. One of these methods is Ultrasound Localization Microscopy (ULM), which uses microbubbles (∼ 2.5 *µ*m in diameter) injected into the bloodstream to generate super-resolved images of cerebral blood flow [13, 14, 15]. In ULM, ultrasound images are acquired over long acquisition times, typically on the order of 10 minutes, at frame rates of about 1 kHz. In post-processing, localization algorithms estimate the position of individual microbubbles with sub-pixel precision, and tracking algorithms reconstruct their trajectories over time. The accumulation of these trajectories yields detailed images of the microvascular network with spatial resolution up to ten times higher than that of conventional power Doppler imaging [13, 15]. More recently, ULM has been extended to dynamic studies of cerebral blood flow, including functional ultrasound localization microscopy (fULM) [16] and vascular pulsatility imaging [17]. However, ULM also has important limitations, including long acquisition times, high computational cost, and the need for microbubble injection. Although microbubbles are generally considered safe, their use remains invasive, may not be suitable in all experimental or clinical settings and is currently approved for only limited clinical applications.

To overcome these limitations, new approaches that avoid the use of contrast-agent microbubbles are being explored. Super-resolution imaging using erythrocytes (SURE imaging) is one such method. It does not require microbubble injection and has been shown to achieve a spatial resolution below the ultrasound wavelength in the rat kidney. Its application to brain imaging is currently under investigation [18, 19].

As a complementary strategy, Null Subtraction Imaging (NSI) is a non-linear beamforming technique that has recently been shown to improve the spatial resolution of power Doppler images without the use of microbubbles [20, 21, 22]. In NSI, three receive apodizations are used: one that produces a zero-mean pattern, creating a sharp null and two with DC offsets, which are combined to generate an artificially sharper main lobe. The DC offset parameter controls the width of this main lobe and, consequently, the degree of spatial-resolution enhancement.

Given these characteristics, the aim of the present work is to evaluate the performance of NSI in functional ultrasound imaging (fNSI) across a range of DC offset values. Specifically, we quantify the resulting trade-off between spatial resolution and SNR under functional imaging conditions and we identify a practical operating range of DC offsets that provides the best compromise between improved spatial localization of the activation maps and reduced temporal fluctuations in the fUS signal.

## 2. Methods

Ultrafast functional ultrasound experiments were performed in vivo in mice during whisker stimulation. RF channel data were acquired using a plane-wave coherent compounding sequence and then processed to obtain power Doppler images. Two processing pipelines were considered: conventional delay-and-sum (DAS) and NSI. Activation maps computed using DAS and NSI were compared using different quantitative metrics.

### 2.1. Animal preparation

All animal experiments and procedures were approved by the local ethics committee [Comisión de Ética en el Uso de Animales, Instituto de Investigaciones Biológicas Clemente Estable (IIBCE), Uruguay; protocol number: 002a/10/2020]. Experiments were conducted in accordance with the relevant regulations (Uruguayan law number 18611) and the ARRIVE guidelines. Experiments were performed in three wild-type C57BL/6 mice (*n* = 3) bred and housed at the IIBCE animal facility under controlled conditions (12 h dark/12 h light cycle, 21 *±* 3 °C), with food and water available ad libitum. Mice were weaned at postnatal day 21, sexed, and identified by ear punching. To avoid craniotomy or skull thinning procedures, experiments were performed in 2-month-old male mice.

Anesthesia was induced using a ketamine/xylazine mixture adjusted to the animal’s weight and diluted twofold in sterile saline solution (0.9% NaCl) [23]. An initial injection corresponding to 85% of the required dose was administered, and supplemental doses were provided as needed based on the assessed depth of anesthesia. Anesthetic level was monitored throughout the experiment using three criteria: (i) slight spontaneous whisker movements; (ii) absence of muscular response to paw pressure; and (iii) absence of the corneal reflex. Under these conditions, animals remained anesthetized for approximately 40 minutes, which was sufficient to complete the experimental protocol.

Approximately 10 minutes after injection, the hair on the scalp was first trimmed with an electric razor and then depilatory cream was applied to remove remaining hair. After an additional 10 minutes, the cream was removed and the head skin surface was exposed. The animal was then placed on a heating pad (HP-1M, Physitemp, USA) mounted on the stereotaxic frame. Body temperature was maintained at 37 °C using a heating pad and rectal probe (HP-1M thermocouple, Physitemp, USA) connected to a temperature controller (TCAT-2DF, Physitemp, USA). This temperature regulation was required to preserve a sufficient level of activity to the applied stimulus.

For imaging, the ultrasound probe was first aligned with the coronal plane. The probe position was then adjusted along the anteroposterior axis to target a coronal section located between *−*1.58 mm and *−*1.82 mm from Bregma. This positioning was guided by visually matching hippocampal and cortical anatomical landmarks to the Paxinos & Franklin atlas [24]. This imaging plane includes the primary somatosensory cortex barrel field (S1BF), linked to vibrissae stimulation.

### 2.2. RF acquisition

RF channel data were acquired using a Verasonics Vantage system equipped with a Vermon linear array probe (128 elements, 0.10 mm pitch, 15 MHz center frequency). Signals were acquired at a sampling frequency of 4 times the transmission frequency, with sampling performed only at 0° and 90° phase. In post-processing, a band-pass filter centered around the carrier frequency and fractional bandwidth of 40% was applied to obtain 4 points per wave-length (full sampling). The fully sampled signals were interpolated up to a sampling frequency of 16 times the transmission frequency to have adequate delay accuracy [25]. Imaging was performed over a 5 mm scanning depth. Ultrafast imaging relied on a plane-wave coherent compounding sequence with four steering angles (− 6°, − 2°, 2° and 6°), transmitted sequentially for each compounded frame. For each angle, the acquisition event was repeated 3 times and the signals were hardware averaged to improve SNR. The sequence was repeated to acquire a total of 350 compounded frames at a frame rate of 500 Hz. These 350 frames constitute the input dataset used to compute power Doppler images using either DAS or NSI, as described in the following sections.

### 2.3. Ultrafast power Doppler processing

After RF acquisition, processing steps were performed offline to generate a power Doppler image from the 350 compounded frames. First, an element sensitivity correction (ESC) was applied to the raw RF channel data to compensate for inter-element receive sensitivity variations, which is particularly important in NSI [20]. ESC gain factors were estimated from calibration acquisitions performed in water using a planar reflector and each receive channel was normalized accordingly prior to beamforming [20].

Beamforming was then performed using conventional DAS on each steering angle, followed by coherent compounding of the four angles to obtain one beamformed frame in complex IQ form. The beamforming grid was defined with a resolution of 25 *µ*m in the lateral direction and 12.5 *µ*m in depth, which corresponds to one point per sample of the interpolated signals. From each 1 s block, a time series of 350 compounded IQ frames was obtained. A lowpass filter was applied to each frame over depth to improve SNR, and was then decimated by the same factor used in interpolation, resulting in a final IQ depth resolution of 25 *µ*m. The resulting IQ frames were clutter-filtered using a singular value decomposition (SVD) filter applied to the spatiotemporal IQ matrix. The same SVD cut-offs were used for both DAS and NSI pipelines: singular values 1–60 were removed to suppress tissue-related coherent components, and singular values above 340 were removed to suppress incoherent noise contributions. These cut-off values were selected empirically to balance tissue suppression and noise rejection while preserving microvascular Doppler signals.

The processing then branches into two reconstructions: conventional DAS and NSI, described below.

#### 2.3.1. DAS power Doppler reconstruction

For DAS, the SVD clutter filter was applied to the 350 compounded IQ frames of each block. A power Doppler image was then computed as the temporal integration of the squared magnitude of the filtered IQ sequence:

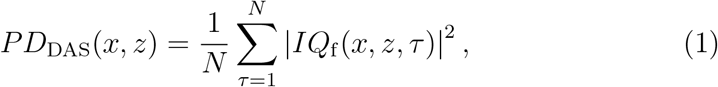

where (*x, z*) are the lateral and axial spatial coordinates, *N* = 350 is the number of compounded frames in the block, *IQ*_f_ denotes the SVD-filtered IQ data and 𝒯*∈* {1, …, *N*} indexes the compounded frames (time).

#### 2.3.2. NSI power Doppler reconstruction

NSI was implemented by applying three receive apodizations to the same RF dataset of each frame: (i) a zero-mean (ZM) apodization with half the aperture weighted +1 and the other half *−*1, (ii) a DC-offset version of the ZM apodization (DC1), and (iii) a laterally flipped version of DC1 (DC2) [21, 20]. The DC offset, denoted *DC*, was varied throughout this study in the range 0.1 to 1 with a step of 0.1. For each frame and each apodization, DAS beamforming was performed (with the same interpolation grid, 25 *µ*m pixel spacing) to obtain three IQ frames: *IQ*_ZM_, *IQ*_DC1_ and *IQ*_DC2_.

Because the NSI subtraction is performed on envelope data, conventional SVD clutter filtering is not directly applicable after NSI subtraction [20]. Following Kou et al., we therefore applied SVD clutter filtering to the beam-formed IQ data prior to envelope detection by forming an augmented spatiotemporal matrix in which, for each time index, the three IQ frames corresponding to ZM, DC1 and DC2 were concatenated and treated jointly [20]. The same SVD cut-offs as in the DAS pipeline were used. After filtering, the three IQ sequences were separated again to yield *IQ*_ZM,f_, *IQ*_DC1,f_, and *IQ*_DC2,f_ .

Envelope images were then computed for each filtered IQ frame, and the NSI image was obtained by subtracting the ZM envelope from the average of the two DC-offset envelopes [21, 20]:

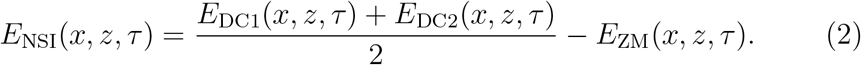

Finally, an NSI power Doppler image was computed from the NSI envelope time series using the same temporal integration as in DAS:

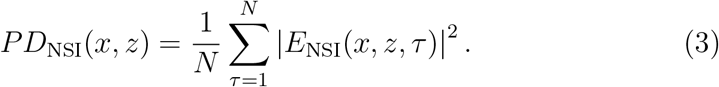

### 2.4. Functional ultrasound experiment and activation maps

Functional ultrasound (fUS) imaging was performed while applying an alternating ON/OFF stimulation pattern to the anesthetized mouse. Whisker stimulation was delivered manually by deflecting the vibrissae with a cotton swab. The stimulation protocol started with a 60 s OFF baseline period, followed by four cycles of 40 s ON and 40 s OFF. Power Doppler images were acquired continuously throughout the protocol, yielding a total recording duration of 380 s with a temporal sampling of one power Doppler image per second. To reduce variability across trials, whisker stimulation was systematically performed by the same researcher.

To quantify stimulus-induced hemodynamic fluctuations, we computed the relative change in cerebral blood volume (CBV) from the power Doppler intensity at each pixel as [23]:

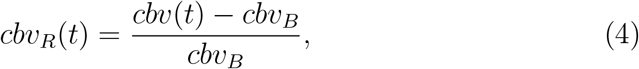

where cbv(*t*) denotes the power Doppler intensity at time *t*, and cbv_*B*_ is its mean value over the first 50 s of the initial 60 s OFF baseline period (absence of stimulation).

Activation maps were constructed by correlating, for each pixel, the temporal CBV signal with the stimulation sequence [23]. Specifically, we defined a binary stimulation regressor *s*(*t*) that takes the value 1 during ON periods and 0 during OFF periods, sampled at the same temporal resolution as the fUS acquisition (1 power Doppler image per second). For each pixel (*x, z*), we computed the Pearson correlation coefficient *r*(*x, z*) over the full recording duration. Statistical significance of the correlation was evaluated using the corresponding *p*-value: correlations with *p >* 0.001 were considered non-significant and discarded when generating thresholded activation maps [23]. In addition, for visualization purposes, we also report unthresholded correlation maps (raw *r*) in selected figures.

### 2.5. NSI vs. DAS evaluation

To compare the performance of NSI and DAS, we evaluated both the static power Doppler images (non-functional) and the functional activation maps and CBV signals obtained during vibrissae stimulation.

For the static power Doppler images, two metrics were used: the SNR computed directly from the images and the spatial resolution, quantified using the isofrequency curve method described by Kou et al. [20].

For the functional experiment, four complementary metrics were used: the activation area, the mean spatial gradient, the Dice similarity coefficient [26] and and the temporal SNR of the CBV signal. The first three metrics were computed from the raw correlation maps, without applying the statistical *p*-value mask, to avoid threshold biases.

To compare NSI and DAS across animals and DC offsets, each metric was expressed as a relative value, defined as the ratio NSI/DAS. This normalization allows comparison across animals with different absolute values.

#### 2.5.1. Image SNR

The image SNR was computed from a segment acquired at the beginning of each functional ultrasound experiment without any stimuli. The signal level *µ* was defined as the mean power Doppler intensity averaged over the first *N* = 50 power Doppler frames acquired during rest and then spatially averaged over all pixels of the image. The noise level *σ* was estimated as the standard deviation of the mean of these same *N* power Doppler images. The image SNR was then computed as

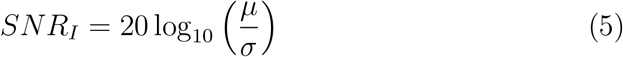

To compare the *SNR*_*I*_ between NSI and DAS across animals and DC offsets, a relative SNR was defined in decibels as Δ*SNR*_*I*_ = *SNR*_*I*_(NSI) *− SNR*_*I*_(DAS). Since SNR is expressed in decibels, this difference directly represents the logarithmic ratio between NSI and DAS signal-to-noise levels. Positive values of Δ*SNR*_*I*_ indicate higher SNR for NSI compared to DAS, whereas negative values indicate lower SNR.

#### 2.5.2. Spatial resolution

Spatial resolution was studied following the isofrequency curve method [27, 20], based on the two-dimensional spatial frequency spectrum of the power Doppler image. After computing the 2-D Fourier transform, we extracted the spectral intensity where the magnitude of the spatial frequency spectrum reaches a fixed wavenumber *k*. For each value of *k*, we computed the mean spectral intensity along this isofrequency contour. For DAS, we identified the spectral intensity at the wavenumber *k*_0_ = 2*π/λ* where *λ* is the wavelength associated with the central frequency (15 MHz). We then located the wavenumber *k*^*′*^ in the NSI spectrum at which the mean spectral intensity equals the DAS mean intensity at *k*_0_. The resolution gain was defined as the ratio *k*^*′*^*/k*_0_, which quantifies how much further into high spatial frequencies the NSI spectrum extends relative to DAS.

#### 2.5.3. Activation area

The activation area was defined as the region enclosed by the contour where the correlation coefficient reaches half of its maximum value.

#### 2.5.4. Mean spatial gradient

To quantify boundary sharpness, we computed the spatial gradient from the activation map using a central difference scheme. The gradient magnitude was calculated at each pixel, producing a map of local changes in the correlation coefficient. We then reported the mean gradient magnitude along the half-maximum contour of the activation map as an indicator of boundary sharpness: steeper gradients correspond to sharper activation edges.

#### 2.5.5. Dice similarity coefficient

The spatial alligment of the activation map with the vascular network was evaluated using the Dice similarity coefficient *DSC* [26], which quantifies the overlap between a binarized activation map (*A*) and a vascular mask (*V*) extracted from the power Doppler image:

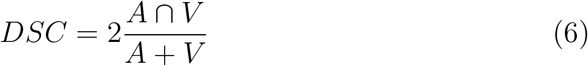

For *A*, pixels with values exceeding half of the maximum correlation value were set to 1. For the vascular mask, rather than using a single global threshold, a locally adaptive threshold was computed for each pixel based on the average intensity within a pixel-centered 3 *×* 3 window, using a sensitivity factor of 0.6 to classify foreground pixels. Dice values closer to one indicate stronger alignment between the activation region and the underlying vasculature.

#### 2.5.6. Temporal signal-to-noise ratio of CBV

To further quantify the SNR ratio of the CBV signal SNR_*cbv*_, we consider time windows extracted from both OFF (absence of stimulation) and ON (stimulation) periods. The *SNR*_*cbv*_ was defined as the ratio between the mean CBV level and its temporal standard deviation within each window:

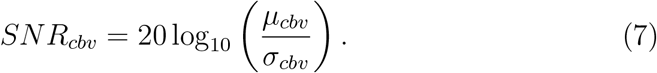

Here, *µ*_*cbv*_ and *σ*_*cbv*_ denote the temporal mean and temporal standard deviation of the CBV signal within the vascular region of interest, computed over the selected window. Higher values of *SNR*_*cbv*_ indicate more stable CBV measurements. A relative temporal SNR was defined in decibels as Δ*SNR*_*cbv*_ = *SNR*_*cbv*_(NSI) *− SNR*_*cbv*_(DAS) to indicate the logarithmic ratio between NSI and DAS signal-to-noise levels.

## 3. Results

Functional ultrasound data were acquired from the three wild-type mice. In the following, one representative mouse is used for qualitative illustration of the reconstruction methods, while all quantitative metrics are systematically reported for the three animals.

### 3.1. NSI vs DAS Evaluation: Non-functional power Doppler images

In this subsection, we compare NSI and DAS using non-functional power Doppler images. We report qualitative comparisons and the quantitative metrics defined in Secs. 2.5.1 (image SNR) and 2.5.2 (spatial resolution).

Figure 1 shows representative power Doppler images reconstructed with DAS and NSI for a DC offset of 0.5. In this example, the two images show similar vascular patterns, with NSI exhibiting subtle improvements in vessel delineation and background levels compared with DAS.

**Figure 1:**
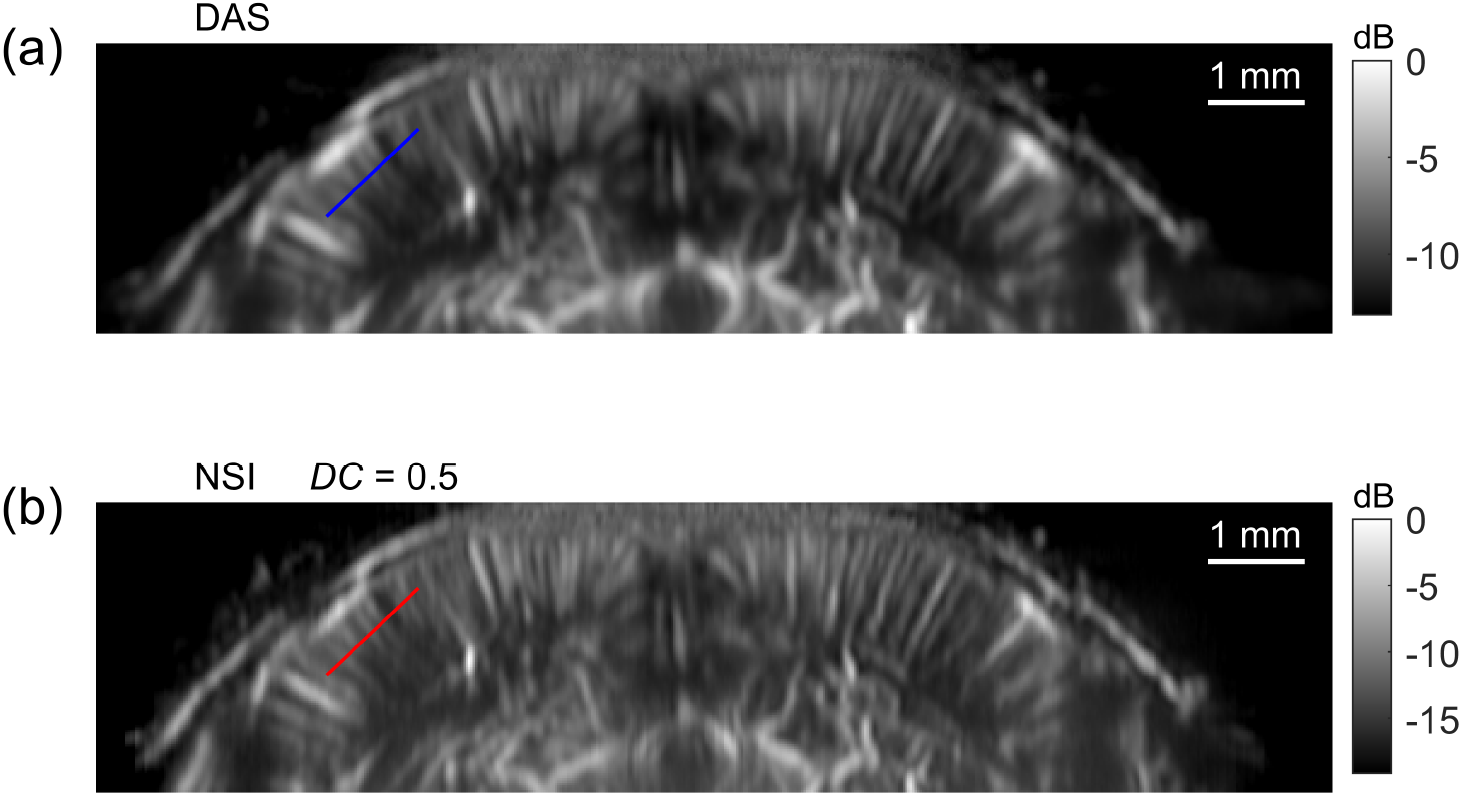
Power Doppler images reconstructed using (a) conventional DAS beamforming and (b) NSI with a DC offset of 0.5. Both images show similar vascular patterns, with NSI exhibiting subtle improvements in vessel delineation and background levels.

A direct comparison between the two methods was performed by extracting intensity profiles in the primary somatosensory cortex along a direction perpendicular to the dominant vessels (Fig. 2). The profile locations are indicated in Fig. 1, with the blue line corresponding to DAS and the red line to NSI. To facilitate comparison, the mean value of each profile was subtracted. The NSI profile shows higher peak amplitudes and a lower background level than DAS, indicating superior contrast between vascular and non-vascular regions. The peaks in the NSI profile are also narrower, which qualitatively reflects a gain in spatial resolution.

**Figure 2:**
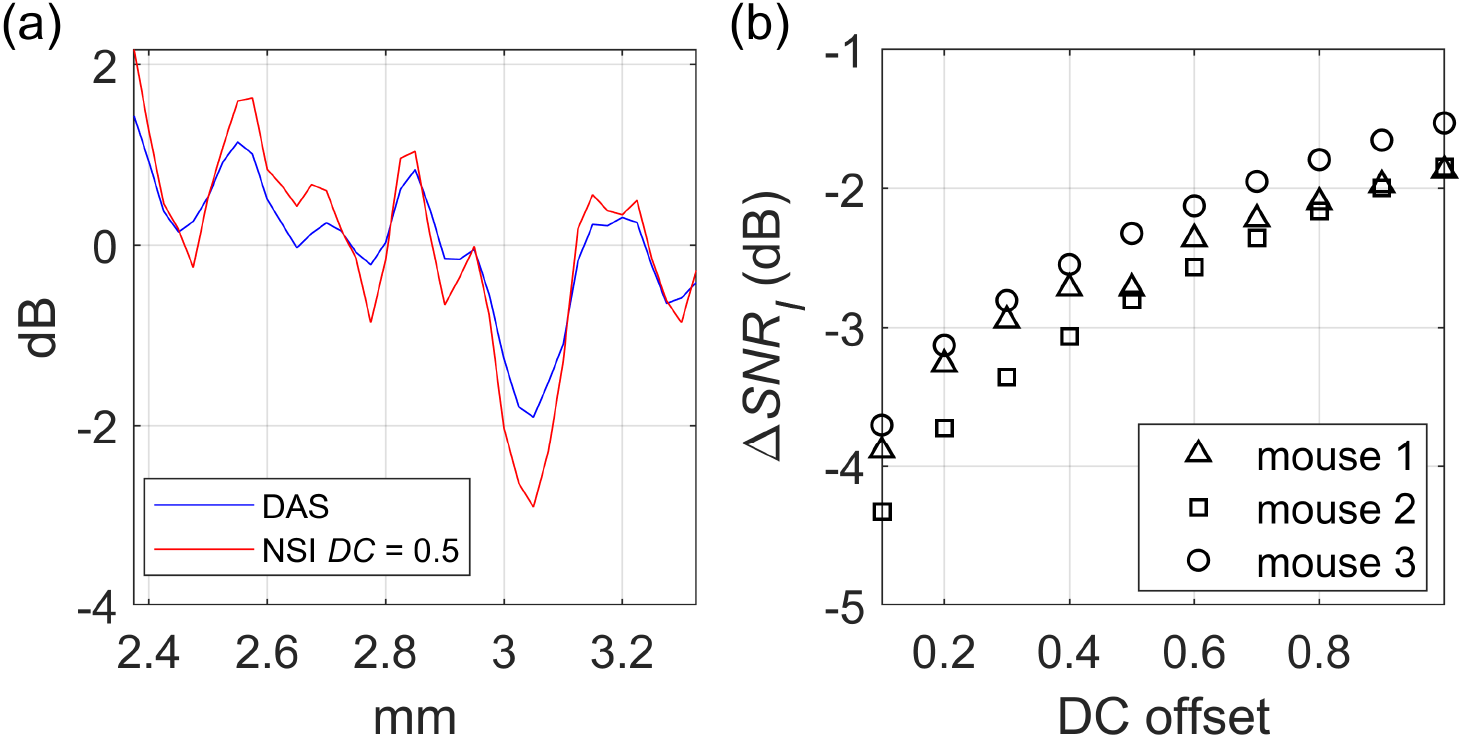
(a) Intensity profiles extracted from the primary somatosensory cortex, perpendicular to the dominant vessels, for DAS (blue) and NSI with *DC* = 0.5 (red). The mean value of each profile was subtracted to facilitate comparison. NSI exhibits higher peak amplitudes, lower background level, and narrower peaks than DAS. (b) Relative image SNR difference, Δ*SNR*_*I*_ as a function of the DC offset for three mice. Triangles, squares and circles correspond to mouse 1, mouse 2 and mouse 3, respectively. Mouse 1 (triangles) corresponds to the representative example shown in (a).

The relative image SNR, computed as described in Sec. 2.5.1, is shown in Fig. 2(b) for three mice, identified by different symbols (triangles, squares and circles). Triangles correspond to the representative mouse shown in Fig. 2(a). The relative image SNR difference, Δ*SNR*_*I*_, ranged from -4.3 dB at *DC* = 0.1 (mouse 2) to -1.5 dB at *DC* = 1 (mouse 3). Thus, NSI consistently exhibited lower image SNR than DAS, although this difference decreased at higher DC offset values.

To further evaluate spatial resolution, we analyzed the two-dimensional spatial frequency spectra of the power Doppler images. Figures 3(a) and 3(b) show the 2-D Fourier transforms for DAS and NSI (*DC* = 0.5), respectively, for the representative mouse. The NSI spectrum exhibits a lateral expansion relative to DAS, revealing a higher contribution of high lateral spatial frequencies, consistent with enhanced lateral resolution. A quantitative metric was obtained from the isofrequency curve analysis described in Sec. 2.5.2, and the corresponding resolution gain is shown in Fig. 3(c) as a function of the DC offset for the three mice. The resolution gain provided by NSI decreases with DC offset, from values around 2 at lower DC offset to values between 1.2 and 1.35 at *DC* = 1, reflecting the trade-off between spatial resolution and SNR.

**Figure 3:**
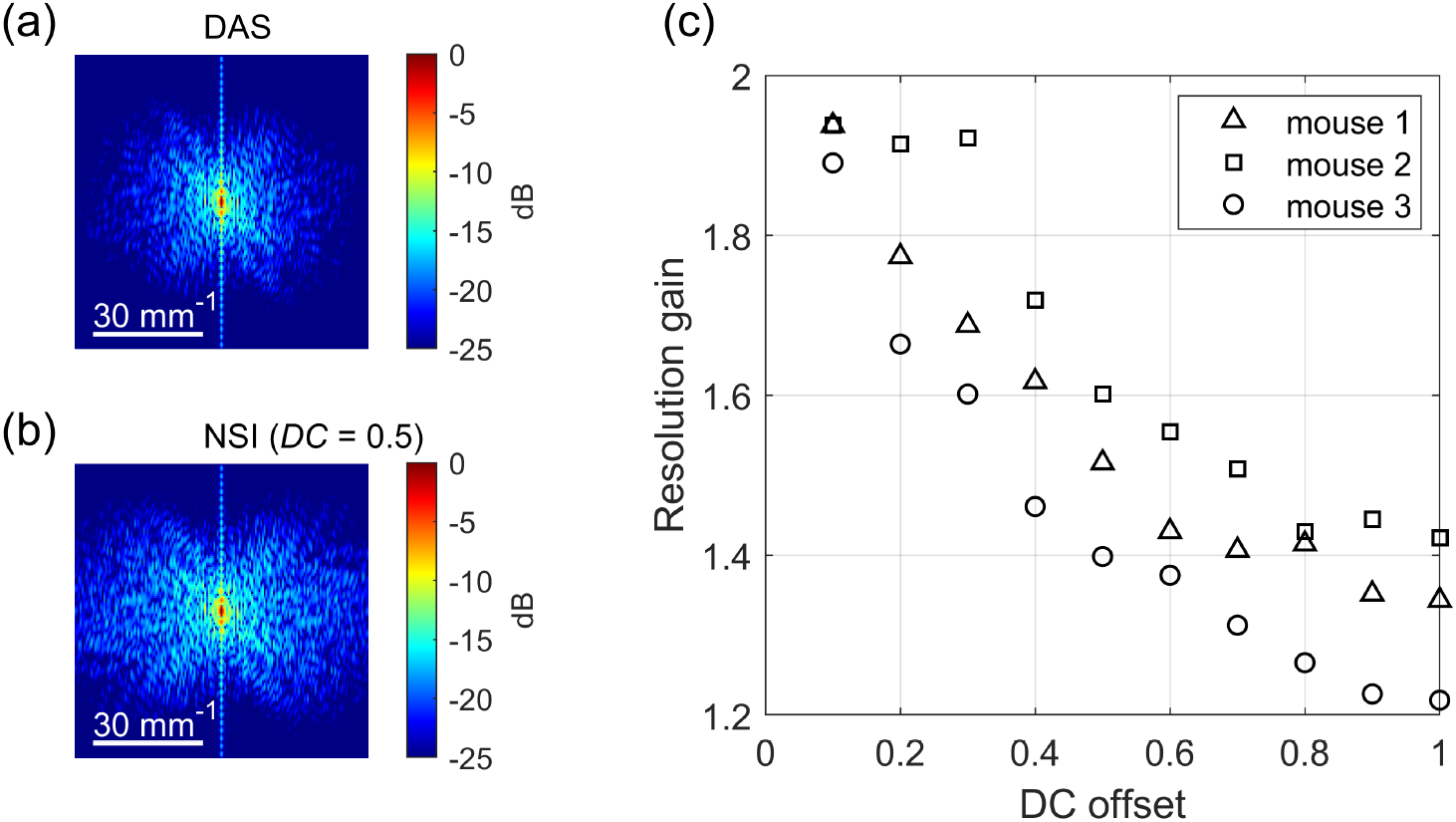
Two-dimensional spatial frequency spectra of the power Doppler images reconstructed using (a) DAS and (b) NSI (*DC* = 0.5) beamforming for the representative mouse. The NSI spectrum exhibits an extended lateral frequency content compared to DAS, indicating enhanced lateral spatial resolution. (c) Resolution gain obtained from the isofrequency curve analysis as a function of the DC offset for the three mice. Triangles, squares and circles correspond to mouse 1, mouse 2 and mouse 3, respectively. Mouse 1 is the representative example shown in (a) and (b).

Overall, these results reveal a clear trade-off between spatial resolution and SNR. Low DC offsets yield better resolution enhancement but reduced SNR, whereas high DC offsets improve SNR at the expense of spatial res-olution. In the following subsection, we characterize how this compromise affects functional ultrasound imaging and identify the most suitable NSI configuration for our imaging setup.

### 3.2. NSI vs. DAS Evaluation: Functional ultrasound experiment

In this subsection, we evaluate how the trade-off between SNR and resolution enhancement introduced by NSI impacts functional activation mapping during periodic vibrissae stimulation. The stimulation pattern and activation map computation are described in Sec. 2.4. Quantitative comparisons are reported using the metrics defined in Secs. 2.5.3 to 2.5.6: activation area (Sec. 2.5.3), mean spatial gradient (Sec. 2.5.4), Dice similarity coefficient (Sec. 2.5.5) and temporal SNR of CBV (Sec. 2.5.6).

Figure 4 shows representative activation maps superimposed on the corresponding power Doppler images obtained with DAS [Fig. 4(a)] and NSI using three different DC offsets: *DC* = 0.2, *DC* = 0.5 and *DC* = 0.8 [Figs. 4(b), 4(c) and 4(d), respectively]. At low DC values (*DC* = 0.2), the activation pattern appears fragmented and spatially fluctuating, which is attributed to the increased noise level introduced by the NSI reconstruction. As the DC offset increases to intermediate values (*DC* = 0.5), the activation region becomes clearly localized, with sharper boundaries and reduced spatial spreading compared with DAS. For higher DC values (*DC* = 0.8), the activation pattern remains visible but appears more spatially extended and diffuse. This behavior reflects the progressive reduction of the spatial-resolution gain as the DC offset increases. The maximum correlation coefficient, however, is slightly higher for DAS than for NSI. This reduction may be associated with the increased noise level observed in NSI power Doppler intensity, particularly at low DC offsets. This aspect will be further addressed in Section 4.

**Figure 4:**
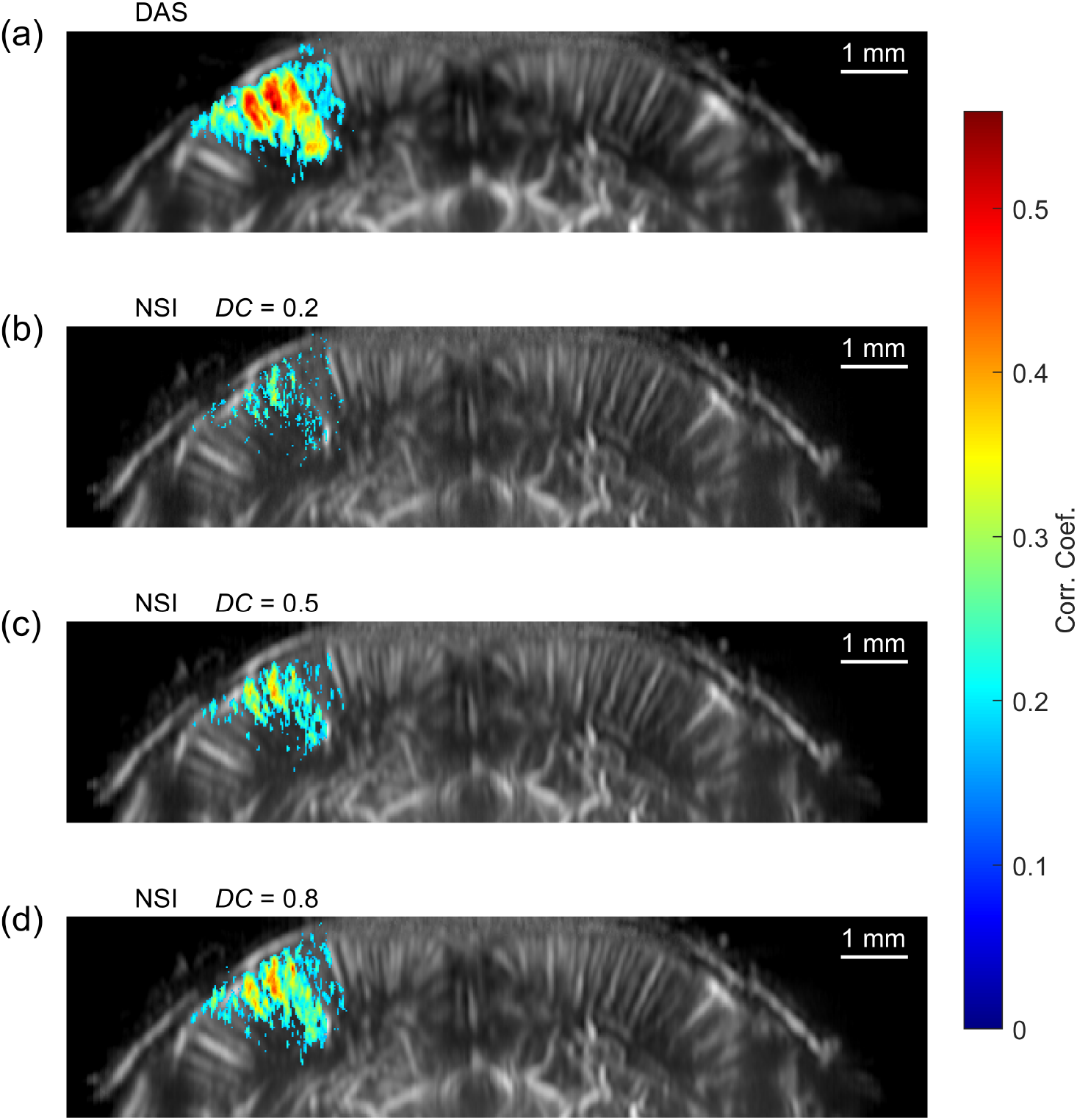
Functional activation maps superimposed on the corresponding power Doppler images for a representative mouse reconstructed using (a) DAS and NSI with three DC offsets: (b) *DC* = 0.2, (c) *DC* = 0.5 and (d) *DC* = 0.8. At low DC values (*DC* = 0.2), the activation map becomes fragmented due to increased noise in the NSI reconstruction. At intermediate DC (*DC* = 0.5), the activation region appears well localized with clearly defined boundaries. At higher DC values (*DC* = 0.8), the activation becomes more spatially extended and diffuse, approaching the appearance obtained with DAS.

In Fig. 5 the activation area is evaluated. Figures 5(a) and 5(b) show the correlation maps for DAS and NSI (*DC* = 0.5), respectively. To avoid interpretation biases associated with applying the statistical *p*-value mask, the colormaps are displayed without masking. The black contour in each panel represents the region above half of the maximum correlation value, providing a consistent criterion within the map for quantifying the activation area. We then compute the relative activated area, defined as the ratio between the NSI and DAS activation areas.

**Figure 5:**
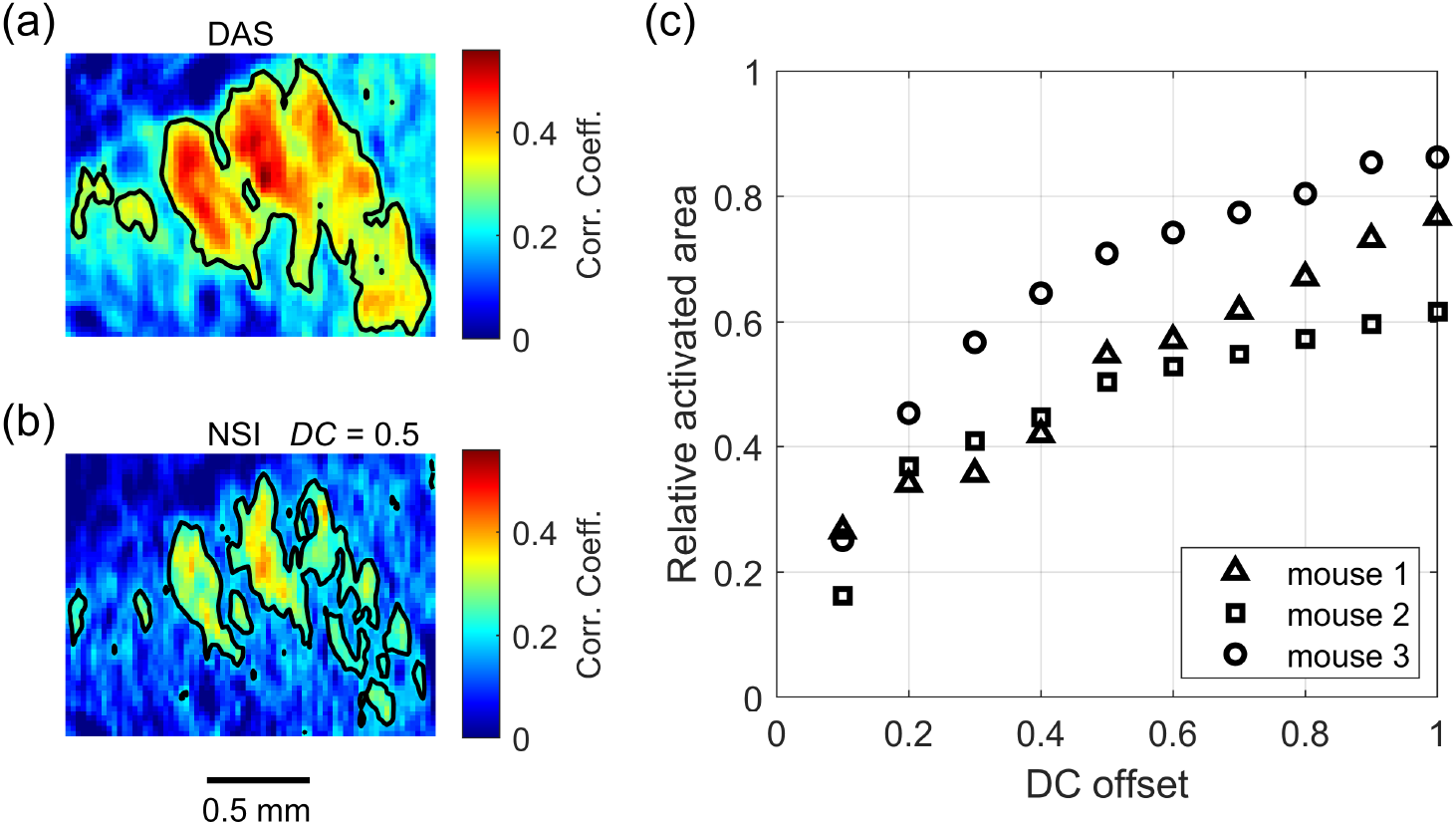
Functional activation maps and activated area analysis. (a) Correlation map obtained with DAS beamforming and (b) correlation map obtained with NSI for a DC offset of 0.5. In both panels, the black contour indicates the region where the correlation coefficient reaches half of its maximum value, which defines the activation area used for quantification. (c) Relative activated area as a function of the DC offset for the three mice. Triangles, squares and circles correspond to mouse 1, mouse 2, and mouse 3, respectively. Mouse 1 (triangles) corresponds to the representative example shown in (a) and (b). Values below unity indicate a reduction of the activated area with NSI compared with DAS.

Figure 5(c) shows the relative activated area for the three mice. For all DC offsets, the relative area is consistently below unity, indicating that NSI yields a smaller activation region compared with DAS. This reduction is consistent with the improved spatial resolution observed in the NSI images. However, a smaller activation area may also arise from reduced SNR, particularly at lower DC values. Indeed, when comparing the relative area with the SNR trends shown in Fig. 2(b), lower activation areas tend to coincide with lower SNR. These observations show that activation area by itself is insufficient to distinguish whether a smaller region comes from better resolution or from increased noise, so additional evaluation is required.

Figures 6(a) and 6(b) show the magnitude of the spatial gradient of the activation map for DAS and NSI with a DC offset of 0.5. The black contour corresponds to the boundary at half of the maximum correlation value, and the mean gradient along this contour (as defined in Sec. 2.5.4) is used as a quantitative measure of boundary sharpness.

**Figure 6:**
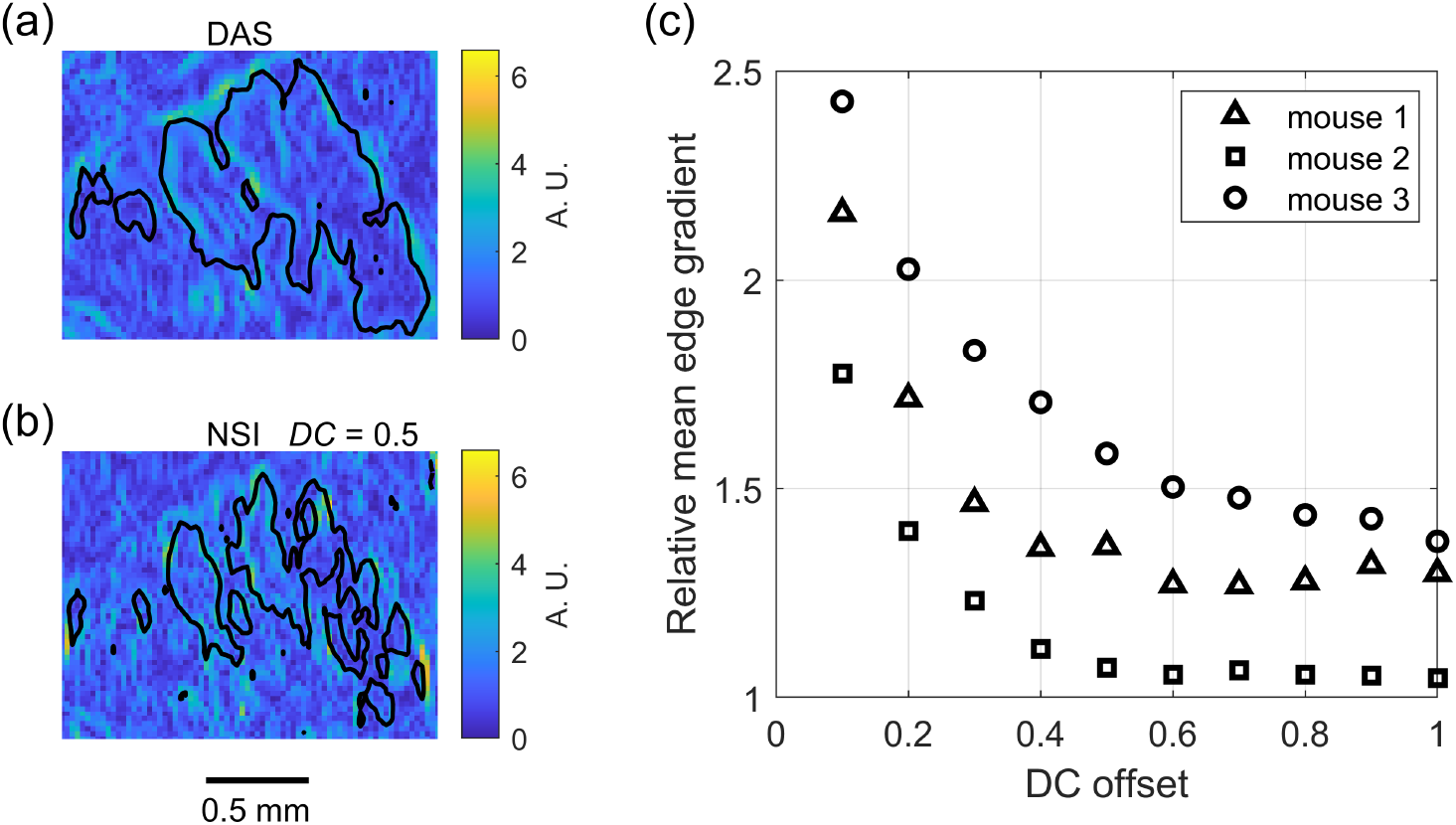
Spatial gradient analysis of the functional activation maps. (a) Magnitude of the spatial gradient of the activation map obtained with DAS and (b) NSI for a DC offset of 0.5. The black contour in each panel indicates the region where the correlation coefficient reaches half of its maximum value. (c) Ratio between the mean gradient values obtained with NSI and DAS, averaged along the half-maximum contour, as a function of the DC offset for the three mice. Values above unity indicate sharper activation boundaries with NSI compared with DAS. Triangles, squares and circles correspond to mouse 1, mouse 2 and mouse 3, respectively. Mouse 1 (triangles) corresponds to the representative example shown in (a) and (b).

The reconstruction methods are compared through the NSI-to-DAS gradient ratio, averaged along the contour for each mouse and plotted as a function of DC offset [Fig. 6(c)]. For all DC values, the ratio is greater than unity, indicating that NSI produces sharper activation boundaries than DAS. Sharper boundaries correspond to more abrupt spatial transitions, which is consistent with higher spatial resolution in the activation maps. According to this metric, NSI provides systematically sharper boundaries than DAS.

The curve also exhibits a change in slope around *DC ≈* 0.3. This feature may reflect the increased noise levels observed at low DC offsets, which can artificially elevate gradient values. For *DC >* 0.5, the ratio is more stable for all animals, reinforcing that NSI yields sharper activation boundaries within this DC range.

The final metric used to evaluate the activation maps is the DSC, defined in Sec. 2.5.5, which quantifies how well the activation region aligns with the underlying vascular structure. Figures 7(a) and 7(b) show the power Doppler image together with the activation contour (in red). The relative DSC was computed for each animal and each DC offset [Fig. 7(c)]. For the mice shown, this ratio is greater than unity for DC offsets above 0.4, indicating that NSI yields activation regions that overlap better with the vascular structures than those obtained with DAS. This suggests that NSI may provide better vascular alignment of the activation map.

**Figure 7:**
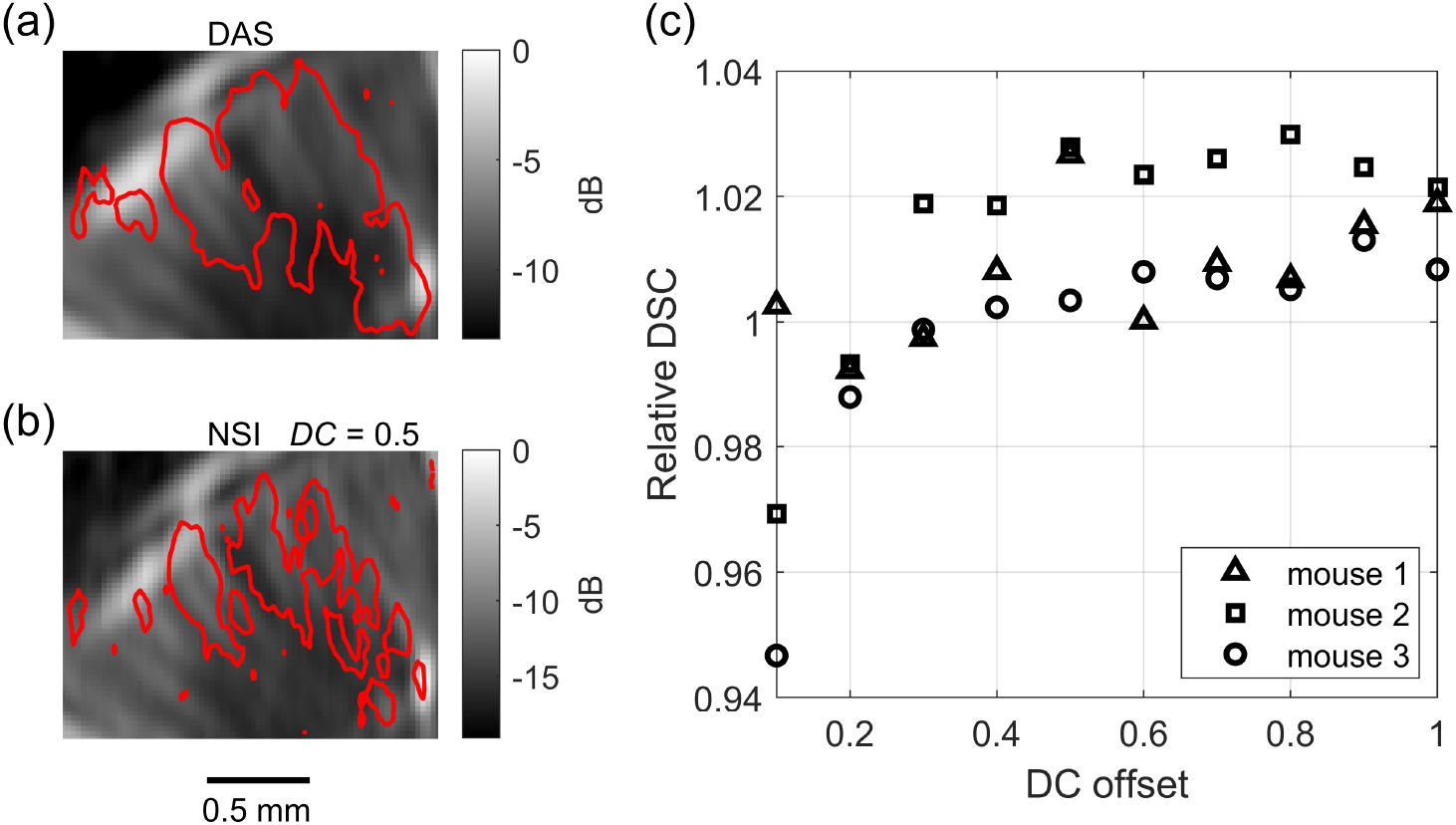
Vascular alignment analysis using the Dice similarity coefficient. (a) Power Doppler image with the activation contour (in red), defined at half of the maximum correlation value, obtained with DAS. (b) Same representation for NSI with a DC offset of 0.5. (c) Relative DSC (NSI/DAS) as a function of the DC offset for the three mice. Values above unity indicate a stronger spatial overlap between the activation region and the underlying vasculature for NSI compared with DAS. Triangles, squares and circles correspond to mouse 1, mouse 2 and mouse 3, respectively. Mouse 1 (triangles) corresponds to the representative example shown in (a) and (b).

Figure 8 shows representative CBV time traces extracted at the pixel corresponding to the maximum activation for DAS [Fig. 8(a)] and NSI using three DC offsets: *DC* = 0.2, *DC* = 0.5 and *DC* = 0.8 [Figs. 8(b), 8(c) and 8(d), respectively]. The blue square waveform indicates the stimulation cycles. The temporal SNR was computed as defined in Sec. 2.5.6 (Eq. 7), using an OFF segment acquired prior to stimulation (first 20 s) and a time window during stimulation (from 70 s to 90 s). At low DC values (*DC* = 0.2), the CBV signal exhibits strong temporal fluctuations due to increased noise in the NSI reconstruction. As the DC offset increases, these fluctuations progressively decrease and the CBV signal becomes more stable, approaching the behavior observed with DAS.

**Figure 8:**
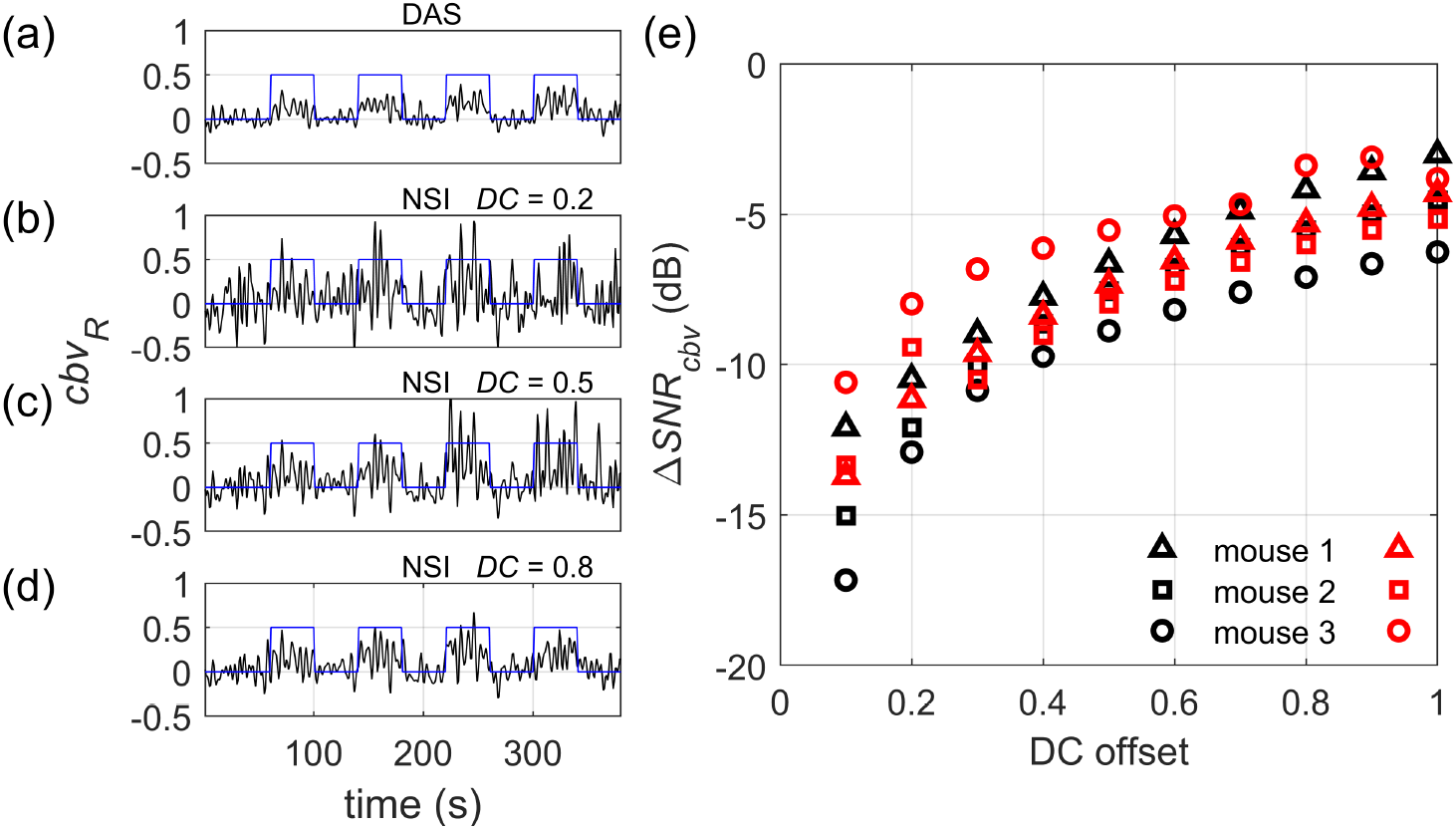
Representative CBV time trace extracted at the pixel corresponding to the maximum activation for (a) DAS beamforming and NSI for (b) *DC* = 0.2, (c) *DC* = 0.5 and (d) *DC* = 0.8. The blue square waveform indicates the stimulation pattern. At *DC* = 0.2, the CBV signal exhibits strong fluctuations. As the DC offset increases, the CBV signal becomes progressively more stable. (e) Temporal SNR difference, Δ*SNR*_*cbv*_, as a function of the DC offset for the three mice, evaluated during the OFF state (black markers) and during activation (red markers). The SNR difference increases with DC offset, indicating improved CBV signal stability for NSI at higher *DC* values.

Figure 8(e) displays the temporal SNR difference, Δ*SNR*_*cbv*_, as a function of the DC offset for the three mice, evaluated both during the OFF state (black markers) and during activation (red markers). For both conditions, the temporal SNR increases with DC offset, with the largest growth occurring between *DC* = 0.1 and *DC* = 0.5. This behavior is consistent across animals. At low DC values, NSI exhibits higher levels of fluctuation due to increased noise, whereas higher DC values lead to more stable CBV signals and improved temporal SNR relative to DAS.

These results further support the existence of a trade-off between spatial resolution and temporal SNR in fNSI. Low DC offsets favor spatial resolution but degrade temporal SNR, whereas intermediate DC values (around *DC ≈* 0.5) provide a better balance between improved spatial localization and sufficient temporal signal stability for reliable functional activation detection.

## 4. Discussion

The analysis of the power Doppler images shows that NSI extends the spatial frequency content toward higher lateral wavenumbers compared with DAS. Resolution gains of up to 2 were observed at low DC offsets, confirming that NSI can effectively enhance lateral resolution. However, this gain pro-gressively decreases as the DC offset increases, approaching DAS-like values at *DC* = 1.

This resolution/noise trade-off is directly reflected in the image SNR, which remained systematically lower for NSI than for DAS, although it increased monotonically with DC offset. A similar trend was observed for the temporal SNR of the CBV signal, where NSI exhibited reduced stability at low DC values and progressively approached DAS performance as DC increased. Taken together, these results indicate that DC tuning is essential to balance spatial resolution and functional signal reliability.

In the functional experiments, NSI produced more spatially confined activation maps than DAS, as reflected by the reduction in activation area for all animals and DC offsets. Although this reduction is consistent with improved spatial localization, it cannot be interpreted as a pure resolution gain, since lower SNR may also contribute to reducing the apparent size of the activated region. This ambiguity highlights the importance of using complementary metrics beyond activation area alone.

The edge-gradient analysis provides additional insight into the spatial properties of the activation maps. For all animals and DC offsets, the NSI-to-DAS gradient ratio remained above unity, indicating that NSI yielded sharper activation boundaries than DAS. Sharper boundaries correspond to more abrupt spatial transitions and are therefore consistent with improved spatial localization in the functional maps. The change in slope observed around *DC ≈* 0.3 may reflect the competing effects of noise amplification at low DC values and genuine resolution enhancement.

The Dice similarity coefficient further suggests that NSI activation maps exhibit better alignment with the vascular structure than those obtained with DAS, particularly for DC offsets above 0.5. This behavior is consistent with the improved vascular alignment expected from enhanced lateral resolution, although this interpretation should remain cautious because the DSC may also be influenced by changes in SNR and map extent.

Finally, the CBV analysis confirms that NSI at low DC values is affected by reduced SNR, which directly impacts temporal stability and, consequently, the peak correlation values. As *DC* increases, *SNR*_*cbv*_ improves, especially above *DC* = 0.5, identifying this region as a practical operating range in which noise and spatial resolution remain reasonably balanced.

This study has some limitations. First, the number of animals was small (*n* = 3), so the present results should be interpreted as an initial proof of concept rather than as a definitive characterization of fNSI performance. Second, whisker stimulation was delivered manually, which may introduce variability despite being systematically performed by the same researcher. Third, the SVD clutter-filter cut-offs were selected empirically for our imaging configuration. Although the same cut-offs were applied consistently to both DAS and NSI pipelines, their optimal values may depend on the experimental setup and should be investigated further in future work.

Overall, our results indicate that NSI can improve the spatial localization of functional ultrasound activation maps relative to DAS, but only within a practical DC range that preserves sufficient image and temporal SNR. In our imaging configuration, intermediate DC values around *DC* ≈ 0.5 provided the most favorable compromise between improved spatial localization and sufficient signal stability.

## 5. Conclusion

In this work, we demonstrated the feasibility of applying NSI to functional ultrasound imaging of the mouse brain. Through a systematic comparison with conventional DAS beamforming across a range of DC offset values, we quantitatively characterized the trade-off between spatial resolution enhancement and SNR inherent to NSI.

NSI yielded more spatially confined activation maps than DAS, as evidenced by the reduction in activation area, the increase in boundary sharpness inferred from the edge-gradient analysis, and the alignment with the vascular network quantified by the Dice similarity coefficient. Together, these findings support improved spatial localization of the functional maps obtained with NSI. However, this spatial improvement was accompanied by a reduction in signal-to-noise ratio at low DC offsets, which can negatively affect the reliability of functional activation maps.

DC offset values around 0.5 were identified as the most practical operating range in our imaging system, where NSI preserved most of its spatial resolution gain while maintaining sufficient temporal SNR for reliable functional detection. This balance is critical for the application of NSI to functional neuroimaging.

Overall, these results establish NSI as a promising strategy for improving spatial localization in functional ultrasound imaging without the need for contrast agents. The quantitative framework presented here provides practical guidance for *DC* parameter selection and supports the use of NSI for high-resolution functional neurovascular imaging.

## Data availability

Data is available upon reasonable request to the corresponding authors.

## Funding Sources

This work was mainly supported by grant CSIC-Grupos I+D 497725-2023-2027. The authors also thank the support of PEDECIBA - Programa de Desarrollo de las Ciencias Básicas, Uruguay and the Agencia Nacional de Investigación e Innovación (ANII), Uruguay. Author also thank SNI-ANII.

## Notes

### Competing Interest Statement

The authors have declared no competing interest.

